# Molecular and cell biological analysis of SwrB in *Bacillus subtilis*

**DOI:** 10.1101/2021.04.30.442225

**Authors:** Andrew M. Phillips, Sandra Sanchez, Tatyana A. Sysoeva, Briana M. Burton, Daniel B. Kearns

**Affiliations:** Department of Biology, Indiana University, Bloomington IN 47408; Department of Microbiology, University of Alabama, Huntsville, AL 35899; Department of Bacteriology, University of Wisconsin Madison, Madison WI 53706

**Author notes:** 1-812-856-2523.

**Keywords:** swarming, type III secretion, T3SS, SwrA, SwrB, motility, flagella, PhoA, FlgW, Flk

## Abstract

Swarming motility is flagellar-mediated movement over a solid surface and Bacillus subtilis cells require an increase in flagellar density to swarm. SwrB is a protein of unknown function required for swarming that is necessary to increase the number of flagellar hooks but not basal bodies. Previous work suggested that SwrB activates flagellar type III secretion but the mechanism by which it might perform this function is unknown. Here we show that SwrB likely acts sub-stoichiometrically as it localizes as puncta at the membrane in numbers fewer than that of flagellar basal bodies. Moreover the action of SwrB is likely transient as puncta of SwrB were not dependent on the presence of the basal bodies and rarely co-localized with flagellar hooks. Random mutagenesis of the SwrB sequence found that a histidine within the transmembrane segment was conditionally required for activity and punctate localization. Finally, three hydrophobic residues that precede a cytoplasmic domain of poor conservation abolished SwrB activity when mutated and caused aberrant migration during electrophoresis. Our data are consistent with a model in which SwrB interacts with the flagellum, changes conformation to activate type III secretion, and departs.

**IMPORTANCE:** Type III secretion systems (T3SS) are elaborate nanomachines that form the core of the bacterial flagellum and injectisome of pathogens. The machines not only secrete proteins like virulence factors but also secrete the structural components for their own assembly. Moroever, proper construction requires complex regulation to ensure that the parts are roughly secreted in the order in which they are assembled. Here we explore a poorly understood activator the flagellar T3SS activation in *Bacillus subtilis* called SwrB. To aid mechanistic understanding, we determine the rules for subcellular punctate localization, the topology with respect to the membrane, and critical residues required for SwrB function.

## INTRODUCTION

Ancestral strains of *Bacillus subtilis* swim in liquid environments and swarm over solid surfaces by synthesizing and rotating flagella (1). Swarming in *B. subtilis* differs from swimming in at least three ways. First, swarming requires the quorum-activated synthesis and secretion of a lipopeptide surfactant to reduce surface tension and create a thin layer of fluid in which to move (2–5). Second, flagella require a greater amount of torque to swarm (6). Third, the rate of *de novo* flagellar synthesis during growth must increase so that the number of flagella on the cell surface exceeds a critical threshold (7–10). Two proteins of poorly understood function, SwrA and SwrB, are required to increase flagellar number. SwrA, is the soluble cytoplasmic master activator of flagellar biosynthesis, which in conjunction with the DNA binding transcription factor DegU, is required to activate the expression of flagellar biosynthesis genes (8,11–15). SwrB, is a single-pass transmembrane protein that increases flagellar number post-transcriptionally (8,16–17). The mechanism of SwrB is poorly understood but genetic evidence suggests that it activates the type III secretion system within the flagellar basal body for the export and assembly of extracellular components.

Flagellar assembly is complex and involves a large number of proteins that must be assembled in a sequential order. The first stage of flagellar assembly is the construction of the basal body in which a protein FliF surrounds and houses a type III protein secretion system (18–20). The basal body is completed by docking of the cytoplasmic C-ring, a cylinder of proteins that contains the rotor for flagellar rotation (21–23). Next, the flagellar type III secretion system exports proteins for the assembly of the axle-like flagellar rod that transits the cell envelope, and the flexible hook that functions as a universal joint (24–26). Once the rod-hook complex achieves a particular length, the secretion system changes specificity and secretes proteins to assemble the long helical flagellar filament (27–32). The completed flagellar structure rotates to act as a propeller and push cells through the environment. In *B. subtilis*, most of the genes required for synthesis of the basal body, rod and hook are encoded by the long, 32 gene, 27 kb *fla/che* operon that is activated by SwrA/DegU (8,13,33–36).

The most distal genes of the *fla/che* operon encode SigD, an alternative sigma factor that directs expression of a regulon that includes genes required for filament assembly, and SwrB, a protein that enhances SigD activity (8,16,35,37–39). To determine how SwrB activates SigD, spontaneous suppressors were isolated that restored swarming to a *swrB* mutant (17). Some suppressor mutations were found in the rotor protein FliG that increased C-ring stability, while other suppressors were shown to increase translation of FliP, a core component of the flagellar type III secretion system (17,19,40–41). It was proposed that SwrB acted as a chaperone to aid assembly of the flagellar basal body which would in turn activate flagellar type III secretion, perhaps by a conformational change propagated through FliF to FliP (17). Accordingly, SwrB-mediated activation of type III secretion would enhance SigD activity by export of its cognate anti-sigma factor antagonist, FlgM (17,42–45). The mechanism by which SwrB promotes basal body assembly or activates the flagellar secretion system is unknown.

Here we explore properties of the SwrB protein to inform the mechanism by which activates type III secretion. We determine SwrB topology in the membrane and find that it is oriented such that the majority of the protein is cytoplasmic. We show that SwrB localizes as puncta at the membrane and that the puncta neither require, nor strictly co-localize with, flagella. Selection for loss-of-function mutations revealed that a charged residue within the transmembrane domain of SwrB was required for punctate localization, and that localization was conditionally required for SwrB activity. Finally, the mutations that most severely impaired SwrB function changed residues at the junction between the transmembrane helix and the large cytoplasmic C-terminus and conferred anomalous mobility in SDS-PAGE. The data are consistent with a model in which SwrB co-localizes with flagella in a manner that is both early and transient, and that a conformational change is likely important for SwrB activity.

## RESULTS

### SwrB localizes as puncta at the membrane independent of flagellar components

SwrB is predicted to be a single-pass transmembrane protein that activates the flagellar type III secretion system, but the mechanism of SwrB is unknown (17). A cell biological approach was undertaken to determine SwrB localization. To generate a functional fluorescent fusion to SwrB protein, we took advantage of the fact that cells mutated for *swrB* were defective for swarming motility (**Fig 1A**) and that swarming can be rescued by complementation when the *swrB* gene was cloned downstream of the *P*_*fla/che*_ promoter and inserted at an ectopic site in the chromosome (**Fig 1B**) (2). Next, an unstructured linker domain and the gene encoding yellow fluorescent protein YFP were fused to the C-terminus of the *swrB* gene within the complementation construct, and the construct was introduced to a cell mutated for *swrB* at the native site. Swarming motility was restored to wild type levels when the SwrB-YFP fluorescent fusion construct was expressed in the *swrB* mutant (**Fig 1B**). We conclude that the SwrB-YFP fluorescent was functional for SwrB activity.

**Figure 1.**
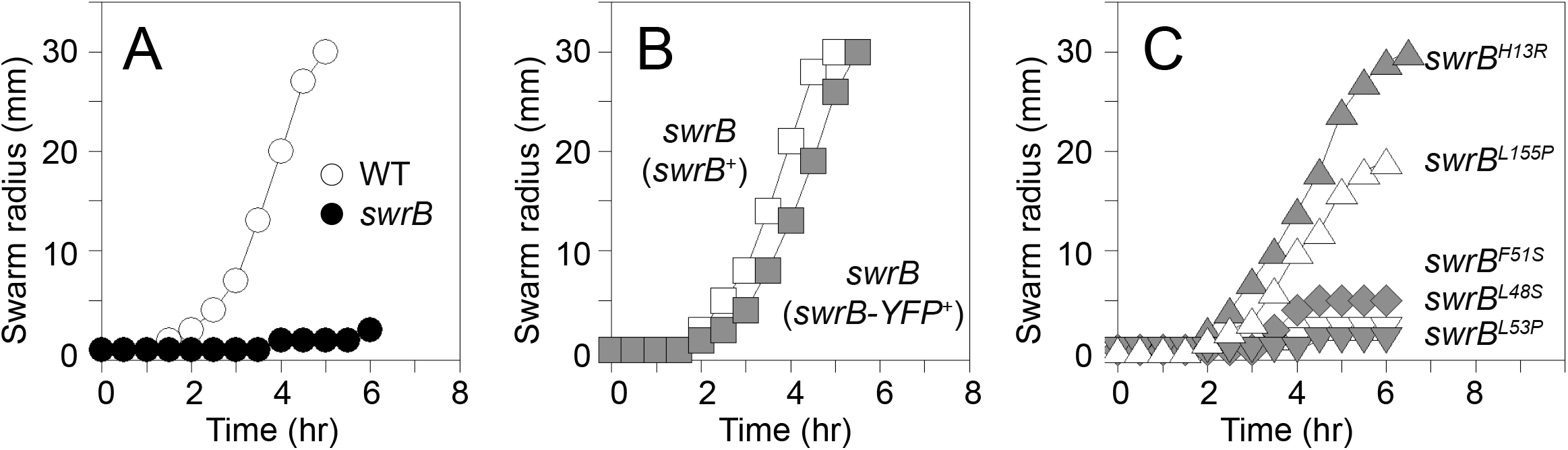
SwrB-YFP is functional and some *swrB* loss-of-function alleles are conditional. Quantitative swarm expansion assays. A) SwrB is required for swarming motility: wild type (open circles, 3610) and *swrB* (closed circles, DS2509); B) SwrB-YPF is functional: *swrB* (*swrB*^*+*^) (open squares, DS2522) and *swrB* (*swrB-YFP*) (gray squares, DS8161); C) Some *swrB* loss-of-function alleles are conditional: *swrB* (*swrB*^*H13R*^) (gray triangles, DS7606), *swrB* (*swrB*^*L155P*^) (open triangles, DS7783), *swrB* (*swrB*^*F51S*^) (gray diamonds, DS7782), *swrB* (*swrB*^*L48S*^) (open inverted triangles, DS7781), and *swrB* (*swrB*^*L53P*^) (gray inverted triangles, DS7669). Genotypes in parentheses indicate complementation constructs. Each line is the average of three replicates.

When observed by fluorescence microscopy, the SwrB-YFP fusion formed faint puncta at the cell membrane reminiscent of the punctate localization of flagellar basal bodies (9) (**Fig 2**). To determine the relative localization of SwrB and flagella, a linker and green fluorescent protein (GFP) was translationally fused to the C-terminus of SwrB at the native *swrB* locus in a strain that also expressed a variant of the flagellar hook protein FlgE (FlgE^T123C^) capable of being labeled with a maleimide reactive-fluorescent dye (46). Fluorescence microscopy indicated that there were fewer SwrB puncta than flagellar hook puncta per cell such that that if SwrB co-localized with flagella, SwrB was only present at a subpopulation of machines (**Fig 3**). Moreover, while some SwrB puncta appeared to co-localize with hook puncta (**Fig 3**, open caret) the majority did not (**Fig 3**, closed caret), suggesting that co-localization was rare. Finally, SwrB puncta were observed in strains mutated for the flagellar basal body baseplate protein FliF (encoded by *fliF*), or mutated for either flagellar type III secretion system protein FlhA or FliP (encoded by *flhA* and *fliP*, respectively) (**Fig 2**). We conclude that SwrB punctate localization is independent of even the earliest components of flagellar assembly (20). We infer that flagellar secretion in a way that either does not require SwrB interaction with the flagellum, or that SwrB co-localizes with flagella at an early step prior to flagellar hook assembly and in a manner that is likely transient.

**Figure 2.**
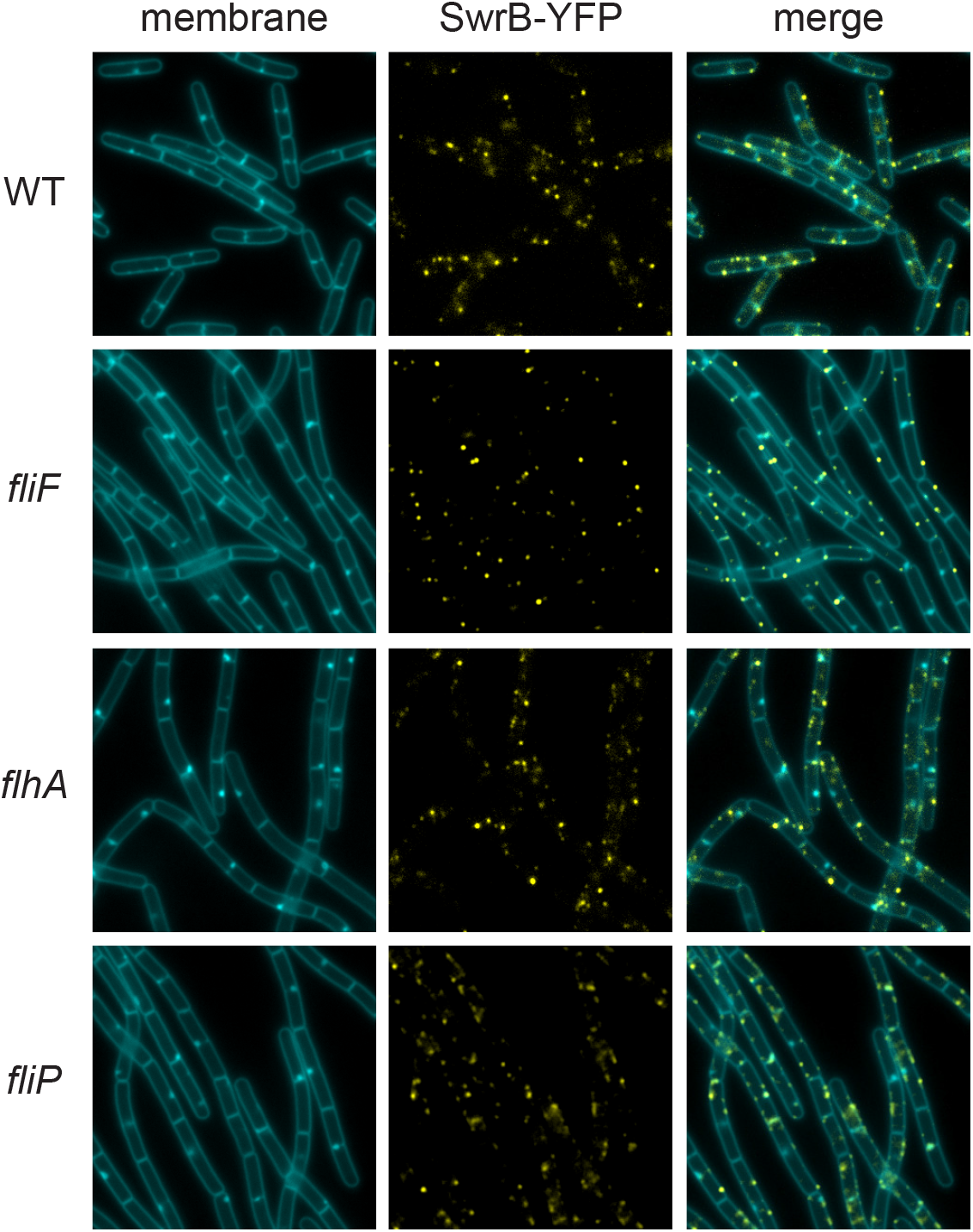
SwrB localizes as puncta that do not require the flagellar basal body. Fluorescence micrographs of YFP translationally fused to SwrB and inserted at the ectopic site in strains carrying a mutated *swrB* gene at the native locus and a mutation in the indicated gene, if any. Membranes were stained with the membrane stain TMA-DPH and false colored cyan. SwrB-YFP was false colored yellow. The following strains were used to generate these panels: wild type (DS8161), *fliF* (DK204), *flhA* (DK310), and *fliP* (DK235).

**Figure 3.**
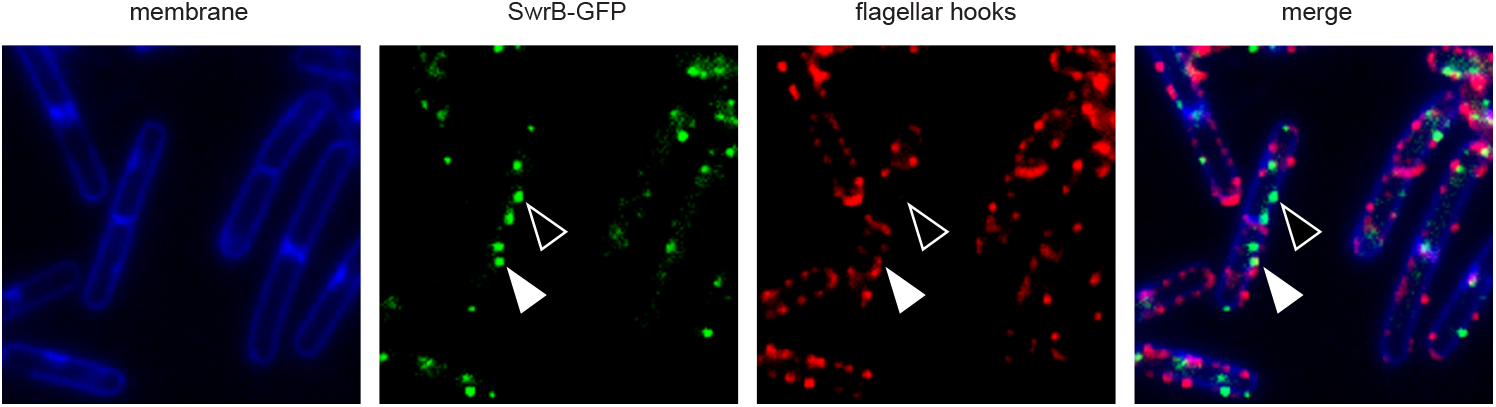
SwrB puncta rarely co-localize with flagellar hooks. Fluorescence micrograph of GFP translationally fused to SwrB and stained for flagellar hooks (FlgE^T123C^) with a fluorescent malemide dye in a wild type background. Membranes were stained with the membrane stain TMA-DPH and false colored blue. SwrB-GFP puncta were false colored green. Hooks were stained with maleimide alexa fluor 592 and false colored red. The following strain was used to generate this panel: wild type (DS9404).

### The large C-terminus of SwrB is intracellular

Primary sequence analysis of SwrB indicates that it is a 167 amino acid protein with a single predicted N-terminal transmembrane domain consistent with a Sec-dependent signal sequence (47–50). Models of SwrB mechanism would be aided by an understanding of its membrane topology. One way of determining membrane topology is by generating fusions of the *E. coli* β-galactosidase LacZ to different regions of the protein of interest. Fusions that display β-galactosidase activity indicate regions of the protein that reside in the cytoplasm because LacZ is only functional in the reducing environment of the cytoplasm (51). To validate topology-dependent LacZ activity in *B. subtilis*, the gene encoding LacZ (*lacZ*) from *E. coli* was codon optimized for *B. subtilis* and expressed from an IPTG-inducible promoter, either in the cytoplasm (no™-LacZ) or in the extracellular environment by fusion to the N-terminal transmembrane segment of the *B. subtilis* flagellar stator protein MotB (MotB™-LacZ) (52–55). Colonies inoculated on plates containing IPTG and the chromogenic substrate 5-bromo-4-chloro-3-inolyl-β-D-galactopyranoside (X-gal) were blue when expressing LacZ in the cytoplasm (**Fig 4A**). By contrast, the colonies were pale when expressing the fusion to MotB, a reduced activity consistent with the known extracellular topology of MotB C-terminal domain and a failure of LacZ to function extracellularly (**Fig 4A**). Finally, sxpression of a fusion of the N-terminal transmembrane segment from SwrB to LacZ (SwrB™-LacZ) produced blue colonies suggesting an opposite orientation to that of MotB (**Fig 4A**). We infer that the C-terminal domain of SwrB is cytoplasmic.

**Figure 4.**
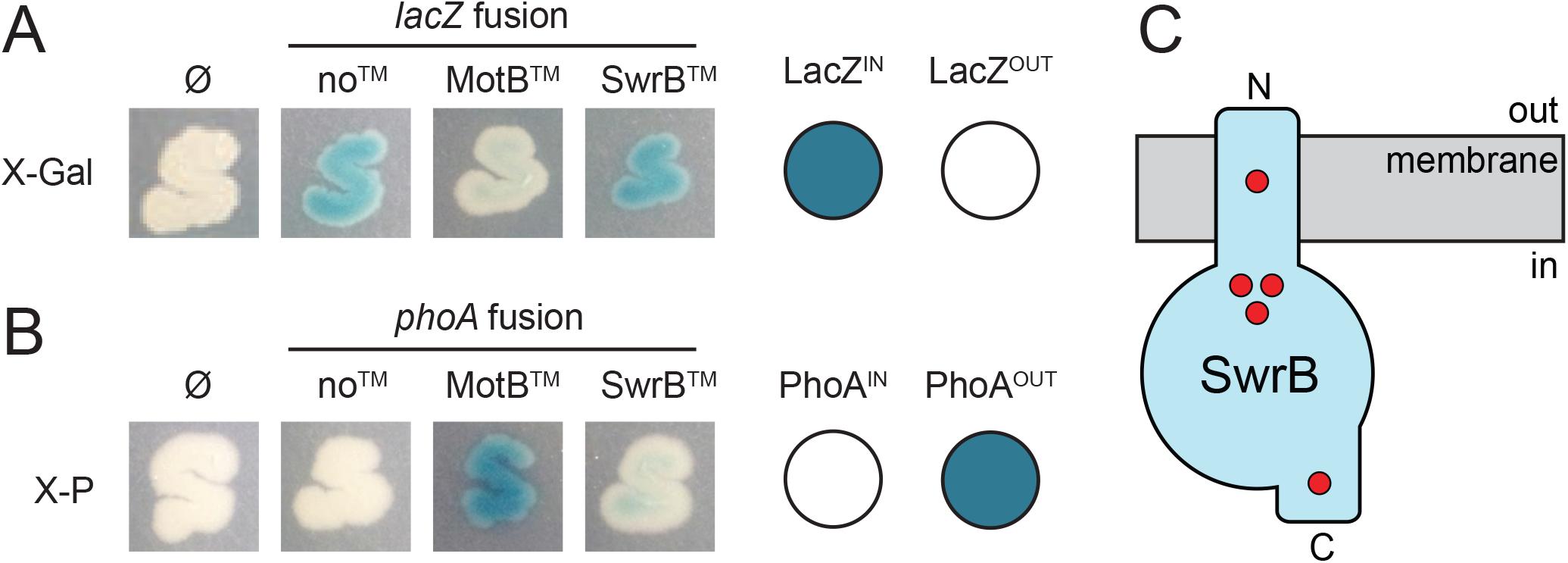
The topology of SwrB positions the C-terminus in the cytoplasm. A) Colonies of strains containing the indicated fusion to *lacZ* on LB containing 1 mM IPTG and the chromogenic substrate X-gal incubated overnight at 30°C. Ø indicates the parental strain that does not contain the *lacZ* gene. The following strains were used to generate the panels: Ø (PY79), noTM (DK9023), motBTM (DK9041), and swrBTM (DK9042). B) Colonies of strains containing the indicated fusion to *phoA* on LB containing 1 mM IPTG and the chromogenic substrate X-P incubated overnight at 30°C. Ø indicates the parental strain that does not contain the *phoA* gene. The following strains were used to generate the panels: Ø (DK3843), noTM (DK8730), motBTM (DK9041), and swrBTM (DK9042). C) Topology prediction of SwrB inserted into the plasma membrane according to results in panels A and B. The putative location of missense mutations outlined in Fig 6A are indicated by red circles.

Another way to determine membrane topology is by generating fusions of the *E. coli* gene *phoA*, encoding the alkaline phosphatase PhoA, to different regions of the protein of interest. Fusions that display alkaline phosphatase activity indicate regions of the protein that extend outside the cytoplasm because PhoA is only functional in the oxidizing external environment (56–57). PhoA fusions are not commonly used directly in *B. subtilis* however, perhaps because wild type colonies have inherent phosphatase activity during vegetative growth that is further induced upon starvation (58–64). Consistent with a basal level of background phosphatase activity, wild type colonies turned blue on media containing the chromogenic alkaline phosphate substrate 5-bromo-4-chloro-3-indolyl-phosphate (X-P). Cells simultaneously mutated for two endogenous alkaline phosphates, PhoA and PhoB (both phylogenetically unrelated to the *E. coli* PhoA protein) (65–66), were white on the same media however, suggesting that *phoA phoB* double mutants would be a good starting background for PhoA fusion analysis (**Fig 4B**). Next, the *phoA* gene from *E. coli* was codon optimized for expression in *B. subtilis*, expressed from an IPTG-inducible promoter, either in the cytoplasm (no™-PhoA) or in the extracellular environment by fusion to the N-terminal transmembrane segment of *B. subtilis* MotB (MotB™-PhoA). In the *phoA phoB* background and the presence of inducer, colonies expressing the no™-PhoA fusion in the cytoplasm were white and colonies expressing the MotB™-PhoA were blue, again consistent with known topology (**Fig 4B**). Expression of a fusion of the N-terminal transmembrane segment from SwrB to PhoA (SwrB™-PhoA) produced colonies that were pale after induction, again suggesting a topology opposite to that of *motB* (**Fig 4B**). We conclude that SwrB is a single-pass transmembrane protein in which the C-terminal domain, comprising the majority of the protein, is cytoplasmic.

### Residues required for SwrB function

To further explore the mechanism of SwrB, a forward genetic approach was undertaken to identify residues required for SwrB function. A screen was devised based on the observation that mutation of *swrB* abolishes σ^D^-dependent gene expression in the absence of the master activator SwrA, and gene expression can be rescued by introduction of a functional *swrB* complementation construct (8,16–17) (**Fig 5**). Next, the *swrB* gene was mutagenized by polymerase chain reaction (PCR) amplification with an error-prone polymerase and the amplicon was cloned downstream of the *P*_*flache*_ promoter between the arms of the *amyE* gene (*amyE::P*_*flache*_*-swrB*^*mut*^). The clones were then pooled and transformed into a *swrA swrB* strain containing a σ^D^-dependent reporter in which the promoter for the *hag* gene (*P*_*hag*_) was transcriptionally fused to the *lacZ* gene encoding β-galactosidase. Finally, the transformants were plated on media containing the chromogenic substrate X-gal such that colonies containing loss-of-function mutations in the *swrB* gene would fail to complement σ^D^-dependent gene expression and confer a white colony phenotype.

**Figure 5.**
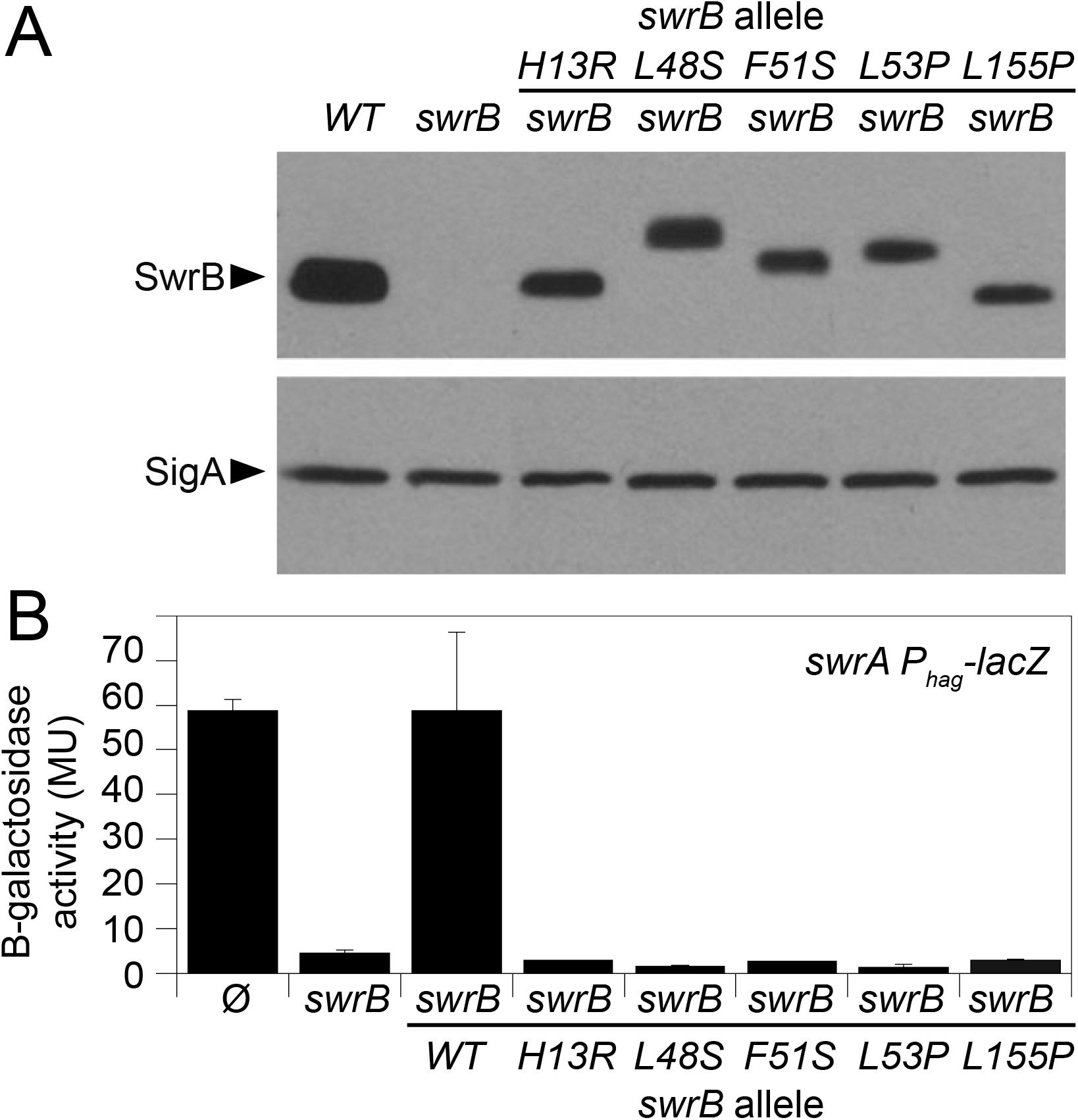
Non-functional alleles of *swrB* that produce stable gene products. A) Western blot analysis of whole cell lysates from the indicated genetic backgrounds probed with anti-SwrB and anti-SigA primary antibodies. Black carets indicate antibody specific targets. Complementation constructs expressing the indicated alleles of *swrB* complements are listed above the black bar. The following strains were used to generate this panel with the complementation construct indicated in parentheses: wild type (3610), *swrB* (DS2509), *swrB* (*swrB*^*H13R*^) (DS7606), *swrB (swrB*^*L48S*^) (DS7781), *swrB (swrB*^*F51S*^) (DS7782), *swrB (swrB*^*L53P*^) (DS7669), *swrB (swrB*^*L155P*^) (DS7783). B) β-galactosidase assays of *swrA P*_*hag*_*-lacZ* transcriptional activity expressed in Miller Units in the indicated genetic backgrounds. The Ø symbol indicates that no further genetic modification was included whereas “*swrB*” indicates the introduction of a *swrB* deletion mutation to the indicated genetic background. Complementation constructs expressing the indicated alleles of *swrB* complements are listed below the black bar. β-galactosidase values presented in **Table S3**. The following strains were used to generate this panel with complementation construct indicated in parentheses: Ø (DS6332), *swrB* (DS6334), *swrB (swrB*^*WT*^) (DS6350), *swrB (swrB*^*H13R*^) (DS7513), *swrB (swrB*^*L48S*^) (DS7443), *swrB (swrB*^*F51S*^) (DS7749), *swrB (swrB*^*L53P*^) (DS7244), *swrB (swrB*^*L155P*^) (DS7750).

Over 4000 transformants were screened and 69 mutants with a white colony phenotype were isolated as SwrB^mut^ alleles. Sequencing of the complementation constructs indicated that many of the mutants contained truncations (nonsense or frameshift mutations), multiple mutations, or defects in the *P*_*flache*_ promoter, and these mutant classes were discarded. Ten alleles contained single missense mutations and were retained. Five of the missense mutations resulted in undetectable levels of SwrB protein by Western blot analysis and were discarded as their defect was likely due to protein instability. The remaining five alleles produced wild type levels of protein (**Fig 5A**) and were defective in rescuing σ^D^-dependent gene expression to a *swrA swrB* mutant background (**Fig 5B**). One mutation was a conservative substitution of a histidine to an arginine (SwrB^H13R^) within the N-terminal transmembrane helix (**Fig 6A, Fig 4C**). Three mutations closely clustered at the junction between the transmembrane segment and the large C-terminal domain (**Fig 6A, Fig 4C**), and each of the junction mutations exhibited anomalous protein mobility when resolved by SDS-PAGE (SwrB^L48S^, SwrB^F51S^, SwrB^L53P^) (**Fig 5A**). The last mutation was near the C-terminus of the protein (SwrB^L155P^) (**Fig 6A, Fig 4C**). We conclude that residues throughout SwrB are required for SwrB function.

**Figure 6.**
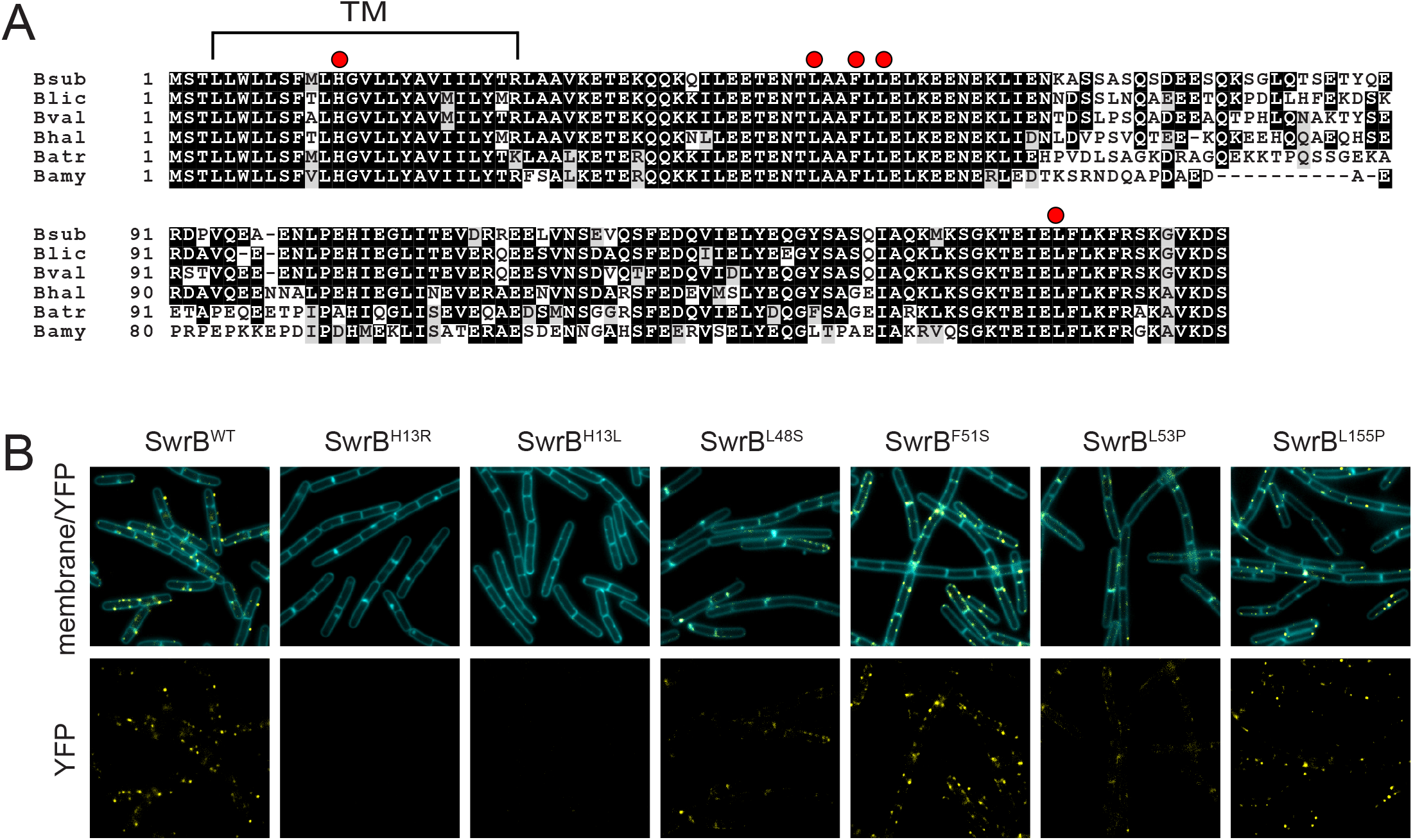
A mutation within the transmembrane domain abolishes punctate localization. A) Multiple-sequence alignment of SwrB from *Bacillus subtilis* (Bsub), *Bacillus licheniformis* (Blic), *Bacillus vallismortis* (Bval), *Bacillus halodurans* (Bhal), *Bacillus atropheus* (Batr), and *Bacillus amyloliquefaciens* (Bamy). Red circles indicate the location of a missense mutation that impairs SwrB function. “TM” and bracket indicate the location of the transmembrane domain. B) Fluorescence micrographs of YFP translationally fused to either the wildtype version of SwrB or SwrB containing the indicated single missense mutations. Membranes were stained with the membrane stain TMA-DPH and false colored cyan. SwrB-YFP false colored yellow. The following strains were used to generate these panels: wild type (DS8161), *swrB swrB*^*H13R*^*-YFP* (DS8364), *swrB swrB*^*H13L*^*-YFP* (DK201), *swrB swrB*^*L48S*^*-YFP* (DS8485), *swrB swrB*^*F51S*^*-YFP* (DS8348), *swrB swrB*^*L53P*^*-YFP* (DS8484), *swrB swrB*^*L155P*^*-YFP* (DS8483).

To determine the consequences of the different alleles of SwrB on localization, the gene encoding YFP was fused to each mutant allele in the complementation construct and introduced to a *swrB* mutant. One substitution, SwrB^H13R^, located within the transmembrane domain resulted in seemingly undetectable SwrB-YFP signal (**Fig. 6B**). SwrB-YFP signal was faint even for the wild type allele, and since SwrB^H13R^ makes similar levels of protein (**Fig 5A**), we suspect that the loss of SwrB^H13R^-YFP signal was due to loss of punctate localization and diffusion of signal throughout the membrane. We note that histidine residues are uncommon within transmembrane domains. When site-directed mutagenesis was used to change histidine 13 to a membrane-compatible hydrophobic leucine residue (SwrB^H13L^-YFP), the resulting construct also abolished punctate localization of SwrB indicating that histidine was required and that the defect was not allele-specific (**Fig. 6B**). The other four loss-of-function alleles, SwrB^L48S^, SwrB^F51S^, SwrB^L53P^, and SwrB^L155P^ all retained punctate localization (**Fig 6B**). We conclude that punctate localization is necessary but not sufficient for SwrB function. We further conclude that punctate localization is mediated by interactions within the transmembrane segment, while the extracellular domain of SwrB may be required for activating the flagellar type III secretion system.

Finally, each of the *swrB* alleles encoded within the ectopic complementation construct were tested for swarming motility in a background mutated for the native copy of *swrB*. Three of the alleles in the junction domain, SwrB^L48S^, SwrB^F51S^, SwrB^L53P^ could not complement the null mutation and exhibited a severe defect in swarming (**Fig 1C**). The SwrB^H13R^ allele in the transmembrane domain and the SwrB^L155P^ allele near the C-terminus however, restored swarming to the *swrB* mutant (**Fig 1C**). We conclude that some of the alleles that were screened for SwrB loss-of-function are instead down-but-not-out allele, as their phenotype was conditional on the simultaneous loss of SwrA. We further conclude that the junction domain of SwrB is the most critical for SwrB activity and confers a loss of function phenotype regardless of genetic background.

## DISCUSSION

SwrB is a narrowly-distributed protein that is required to increase flagellar number during swarming motility. In the absence of SwrB, flagellar assembly is restricted at an unusual step as there is a dramatic reduction in the number of flagellar hooks despite a wild type number of basal bodies (17). Analysis of suppressor mutations that restored swarming to a *swrB* mutant suggested the *B. subtilis* flagellar type III secretion system is assembled in an inactive state, and becomes activated when surrounded by the basal body protein FliF, docked with a functional cytoplasmic C-ring made of FliG, FliM and FliY (17,45). It was proposed that complete assembly of the super-complex acted as a checkpoint so that rod subunits would not be secreted until the basal body, upon which they are to be polymerized, is mature. In the context of this model, SwrB was proposed to be a chaperone that accelerated basal body maturation but the mechanism of how SwrB might serve such a role was unclear. Here we further characterize SwrB and reveal constraints on its putative mechanism.

The localization of SwrB informs function. Fluorescent fusions to SwrB were functional and localized as faint puncta at the cell membrane. The low fluorescence intensity of the fusion indicates that SwrB is in low-abundance, and the number of the SwrB puncta were fewer than the number of basal bodies in the cell. Both the low abundance and substoichiometric ratio of SwrB to its putative target support the notion that SwrB acts catalytically to promote basal body assembly. Moreover, puncta of SwrB formed even in the absence of the basal body and rarely co-localized with flagellar hooks. Thus, SwrB either accelerates basal body maturation remotely or, more likely, localizes to basal bodies transiently prior to rod and hook secretion. Remote activity could include the synthesis of a small molecule which somehow potentiates basal body assembly, or the post-translational modification of a key structural component prior to incorporation into the flagellum. Transient association, on the other hand, would be consistent with a protein-folding chaperone.

SwrB is not homologous to proteins of known function and thus if it is involved in the enzymatic synthesis of a small molecule, or modifies a flagellar structural subunit, it does so using a domain not previously studied. Random mutagenesis was used to determine putative “active site” residues required for SwrB function. A charged residue, histidine^13^, within the transmembrane segment was required for punctate localization, but mutation of that residue acted as a down-but-not-out allele defective only when SwrB levels were low. Thus, puncta formation may be a way to increase local SwrB protein concentration to levels necessary for activity but appears to have reduced importance during conditions that promote swarming motility when SwrB levels rise. Three hydrophobic residues leucine^48^, phenylalanine^51^, and leucine^53^ distal to the transmembrane segment, when mutated, conferred severe functional defects under all conditions tested and substitution of the critical residues resulted in anomalous protein mobility in SDS-PAGE. Anomalous migration has been attributed to differential access of detergent, perhaps by constrained secondary or tertiary folding (67–70), suggesting that domain flexibility and dynamism may be critical for SwrB function. Finally, the three critical residues, and another conditionally-required residue leucine^155^, flank a region of low sequence conservation (**Fig 6A**). The region of low conservation seems inconsistent with a sequence-constrained enzymatic domain, but could be an interaction surface that specifies the SwrB target.

Whether SwrB directly interacts with the flagellar basal body and/or any of the flagellar structural subunits is unknown. The range of potential targets for interaction however is constrained by SwrB topology. SwrB is a single-pass transmembrane protein with an N-terminal transmembrane helix consistent with a Sec-dependent secretion signal. Normally, Sec-secretion signals are inserted such that the N-terminus is oriented towards the cytoplasm guided by positively charged amino acids proximal to the helix according to the “positive inside rule” (50,71). SwrB, however, has positively charged amino acids distal to the helix predicting an atypical topology in which the N-terminus was extracellular. Here we use translational fusions of β-galactosidase and alkaline phosphatase to support the topology prediction and demonstrate that SwrB is oriented such that the majority of its mass is cytoplasmic. Thus, if SwrB interacts with a flagellar component, we predict it would be a component largely contained within the cytoplasm as well. One candidate target could be the FliF-FliG rotor surface as mutations that stabilized FliG assembly suppressed the absence of SwrB. Other suppressors of *swrB* overexpressed the secretion protein FliP and thus SwrB could interact with the cytoplasmic domains of one or more flagellar type III secretion proteins.

With respect to type III secretion, it was brought to our attention that a distant relative of SwrB in *Campylobacter jejuni* called FlgW, interacts with a protein called FliO (personal communication, Beile Gao; 72). FliO is found in flagellar type III secretion systems but not in paralogous injectisomes, and recent work indicates that FliO is a regulator of the flagellar type III protein FliP (73–75). In *B. subtilis*, FliO is predicted to contain two transmembrane segments, and like SwrB, FliO is predicted to have a large C-terminal domain in the cytoplasm. If SwrB were to bind to and regulate FliO, it could explain the increased frequency of flagellar type III secretion activation as well as the observation that excess FliP could bypass the absence of SwrB, perhaps by FliO titration. To be clear, FliO as a target of SwrB is entirely speculative and nearly all research on FliO activity has been performed in *Salmonella enterica*, an organism lacking a SwrB homolog. We note however the uncanny similarities in localization, topology, and function between SwrB and Flk, a poorly understood single-pass transmembrane protein that regulates the *S. enterica* flagellar type III secretion system (76–79). Thus, we wonder whether the coupling of flagellar basal body maturation to secretion activation is conserved even if the proteins that regulate the checkpoint are not.

## MATERIALS AND METHODS

### Strains and growth conditions

*B. subtilis* strains were grown in lysogeny broth (LB) (10 g tryptone, 5 g yeast extract, 5 g NaCl per L) broth or on LB plates fortified with 1.5% Bacto agar at 37°C. When appropriate, antibiotics were included at the following concentrations: 10 µg/ml tetracycline, 100 µg/ml spectinomycin, 5 µg/ml chloramphenicol, 5 µg/ml kanamycin, and 1 µg/ml erythromycin plus 25 µg/ml lincomycin (*mls*). For the swarm expansion assay, swarm agar plates containing 25 ml LB fortified with 0.7% Bacto agar were prepared fresh and the following day were dried for a total of 20 minutes in a laminar flow hood (see below).

### Strain construction

All constructs were first introduced into the domesticated strain PY79 or the cured strain DS2569 by natural competence and then transferred to the 3610 background using SPP1-mediated generalized phage transduction (80–81). All strains used in this study are listed in Table 1. All primers used in this study are listed in **Supplemental Table S1**. All plasmids used in this study are listed in **Supplemental Table S2**.

**Table 1:**
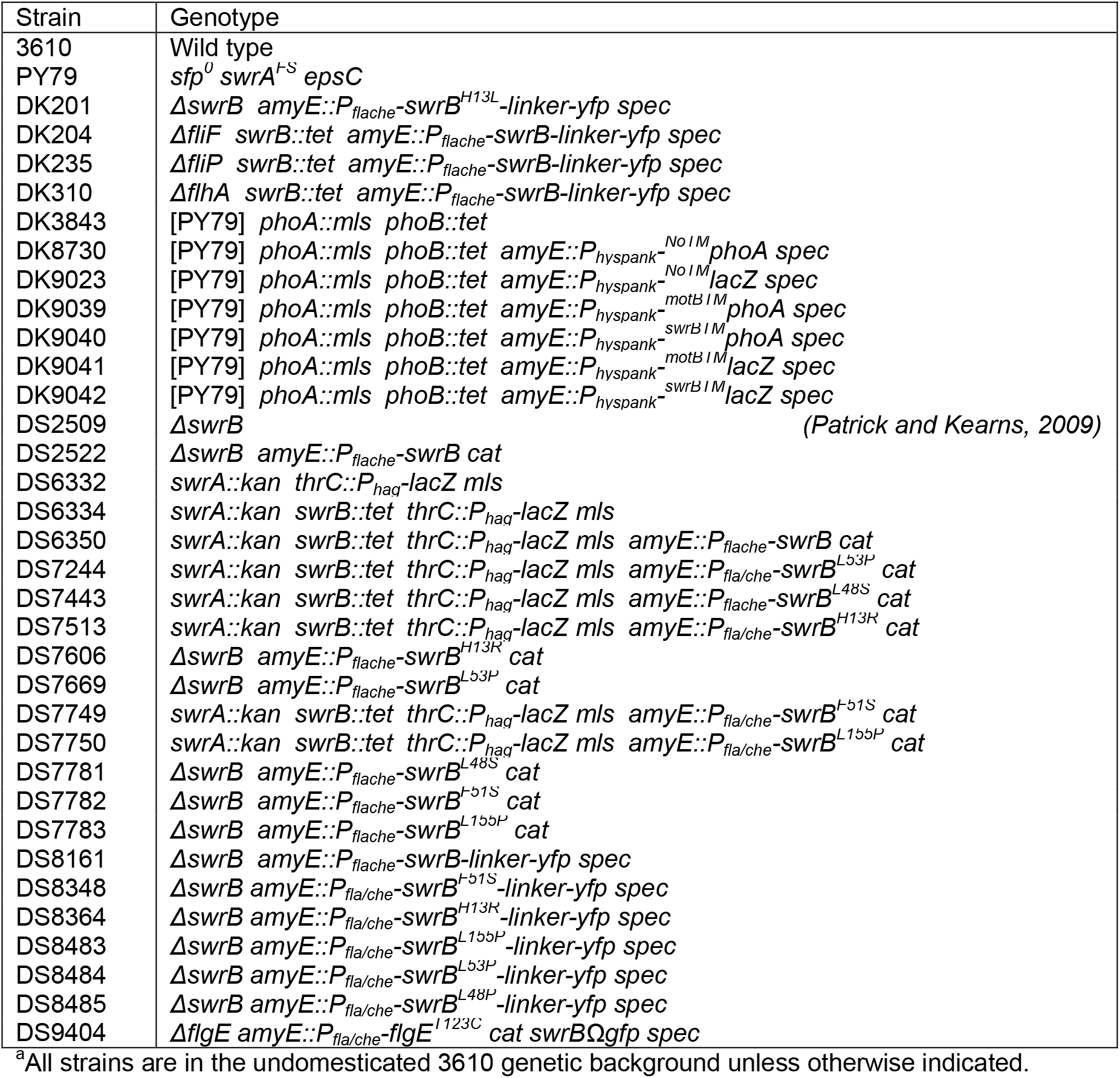
Strains^a^.

#### Fluorescent fusions

To generate the P_*flache*_-*swrB-linker-yfp* translational fusion construct (pAP36), a PCR product containing *linker-yfp* was amplified from *B. subtilis* DS5436 chromosomal DNA using the primer pair 2634/2635 and digested with SalI and BamHI. The fragment was then ligated into the SalI and BamHI sites of pAH25 containing a spectinomycin resistance cassette between two arms of the *amyE* gene (generous gift by Amy Camp, Mount Holyoke College) to generate pAP35. Next, a PCR product containing P_*flache*_-*swrB* was amplified from *B. subtilis* DS6350 chromosomal DNA using the primer pair 2636/2637 and digested with EcoRI and SalI. The fragment was then ligated into the EcoRI and SalI sites of pAP35 to generate pAP36.

To generate the mutated *P*_*flache*_*-swrB-linker-yfp* translational fusions were amplified using primer pair 2636/2637 and chromosomal DNA from the following strains: DS7749 (*swrB*^*F51S*^), DS7513 (*swrB*^*H13R*^), DS7750 (*swrB*^*L155P*^), DS7244 (*swrB*^*L53P*^), and DS7748 (*swrB*^*L48S*^). Each fragment was purified, digested with EcoRI and SalI and ligated into the EcoRI/SalI sites of pAP35 to generate pAP37, pAP40, pAP41 pAP42, and pAP43 respectively.

To generate *swrBΩGFP spec*, a fragment containing *swrB* was amplified from *B. subtilis* 3610 chromosomal DNA using the primer pair 384/385 and digested with EcoRI and XhoI, and the fragment containing *gfp* was amplified from pMF35 using the primer pair 995/996 and digested with XhoI and HindIII (82). The two fragments were then simultaneously ligated into the EcoRI and HindIII sites of pUS19 to generate pDG136 (83).

#### PhoA fusion topology reporters

To generate a *phoA* topology reporter system for *B. subtilis*, the *phoA* gene from *Escherichia coli* was codon optimized for *B. subtilis* by gene synthesis (Integrated DNA technologies) with a 5’ fusion of the *motB* gene from *B. subtilis*. The synthesized gene fragment was then used as a template and PCR amplified with primer pair 7476/7478 and 7477/7478, digested with SalI/SphI and cloned into the SalI/SphI sites of pDR111 containing the *P*_*hyspank*_ promoter, a polylinker, the *lacI* gene, and a gene encoding for spectinomycin resistance between the arms of the *amyE* gene for ectopic integration (generous gift of David Rudner, Harvard Medical School), to generate plasmids pDP545 (*amyE::P*_*hyspank*_*-motB’phoA*) and pDP546 (*amyE::P*_*hyspank*_*-no*^*TM*^*phoA*), respectively. The plasmid pDP545 was constructed such that the 5’ end of theoretically any gene could be cloned between the SalI site and an internal NheI site immediately upstream of the *phoA* coding region to generate in-frame translational fusions to *phoA*. As such, extended fragments of the 5’ ends of the *motB* and *swrB* genes were PCR amplified using pDP545 and DK1042 chromosomal DNA as a template respectively and and primer pairs 7476/7507 and 7489/7490 respectively. Each product was purified, digested with SalI and NheI, and cloned into the SalI/NheI sites of pDP545 to generate pDP566 (*amyE::*_*Physpank*_*-motB*^*TM*^*phoA*) and pDP567 (*amyE::P*_*hyspank*_*-swrB*^*TM*^*phoA*) respectively

#### LacZ fusion topology reporters

To generate a *lacZ* topology reporter system for *B. subtilis*, the *lacZ* gene from *Escherichia coli* was codon optimized for *B. subtilis* by gene synthesis (Blue Heron Gene Synthesis) and cloned into the HindIII/SphI sites of pDR111 to create pDP559 (*amyE::P*_*hyspank*_*-no*^*TM*^*lacZ*). The plasmid pDP559 was constructed such that the exact same fragment cloned upstream of *phoA* in pDP545 could also be cloned into immediately upstream of the *lacZ* coding region. As such, extended fragments of the 5’ ends of the *motB* and *swrB* genes were PCR amplified using pDP545 and 3610 chromosomal DNA as a template respectively and primer pairs 7476/7507 and 7489/7490 respectively. Each product was purified, digested with SalI and NheI, and cloned into the SalI/NheI sites of pDP559 to generate pDP568 (*amyE::*_*Physpank*_*-motB*^*TM*^*lacZ*) and pDP569 (*amyE::P*_*hyspank*_*-swrB*^*TM*^*lacZ*) respectively.

#### phoA::mls

To generate the *phoA::mls* marker replacement insertion/deletion allele, the region upstream of *phoA* was PCR amplified using primers 7382/7383 and the region downstream of *phoA* was PCR amplified using primers 7384/7385 using chromosomal DNA from DK1042 as a template. Next the *erm* cassette (conferring resistance to erythromycin and lincomycin, *mls*) was amplified using primers 3250/3251 using pAH52 (generous gift from Amy Camp, Mount Hoyloke College) as a template. The three PCR products were purified, mixed in equal ratios in an isothermal assembly reaction. The reaction was then transformed into PY79 selecting for *mls* resistance.

#### phoB::tet

The *phoB::tet* allele was acquired from Patrick Eichenberger, New York University (84).

#### Allelic replacement

The *swrB*^*H13L*^ allele was generated using isothermal “Gibson” assembly (ITA) (85). A fragment containing the 5′ half of *swrB*, the spectinomycin-resistance marker (*spec*), and the 5’ *amyE* arm approximately 3000 bp upstream of the *swrB* gene was PCR amplified from DS8161 using primer pair 3177/3292, and a fragment containing the 3′ half of *swrB*, a polylinker, and the 3’ *amyE* arm approximately 3000 bp downstream of the *swrB* gene was PCR amplified using primer pair 3291/3180. Primers 3292 and 3291 are reverse complements and were used to change codon 13 of *swrB* from CAC (histidine) to CTT (leucine).

Isothermal assembly reaction buffer (5×) (500 mM Tris-HCl [pH 7.5], 50 mM MgCl_2_, 50 mM dithiothreitol [DTT] [Bio-Rad], 31.25 mM polyethylene glycol 8000 [PEG 8000] [Fisher Scientific], 5.02 mM NAD [Sigma-Aldrich], and 1 mM each deoxynucleoside triphosphate [dNTP] [New England BioLabs]) was aliquoted and stored at −80°C. An assembly master mixture was made by combining prepared 5× isothermal assembly reaction buffer (131 mM Tris-HCl, 13.1 mM MgCl_2_, 13.1 mM DTT, 8.21 mM PEG 8000, 1.32 mM NAD, and 0.26 mM each dNTP) with Phusion DNA polymerase (New England BioLabs) (0.033 units/μl), T5 exonuclease diluted 1:5 with 5× reaction buffer (New England BioLabs) (0.01 units/μl), *Taq* DNA ligase (New England BioLabs) (5,328 units/μl), and additional dNTPs (267 μM). The master mix was aliquoted as 15 μl and stored at −80°C.

The two DNA fragments were combined at equimolar amounts to a total volume of 5 μl and added to a 15-μl aliquot of prepared master mix. The reaction mixture was incubated for 60 min at 50°C. The completed reaction generated a 6 kb DNA fragment that was PCR amplified using the primer pair 3177/3180, and then directly transformed into PY79. Chromosomal DNA was purified from colonies resistant for spectinomycin and PCR amplified using primer pair 2636/2635 to determine which isolate had retained the *swrB*^*H13L*^ allele.

### PCR mutagenesis

To generate a pool of *swrB* mutants, primer pair 861/862 was used to amplify the *amyE* construct containing *swrB* open reading frame under the control of the P_*flache*_ promoter (*P*_*flache*_*-swrB cat*). DS6350 chromosomal DNA was used as a template and Expand polymerase with Expand Buffer 2 (Roche). The resulting PCR fragment was purified using the QIAquick PCR purification kit (Qiagen) and transformed into PY79. All of the resulting colonies were pooled together and used to generate a lysate library of *swrB*^*mut*^ complementation constructs. The lysate library was SPP1 subsequently transformed into DS6334, and the resulting transductants were patched onto LB containing X-Gal to screen for *swrB* mutants that fail to complement.

### Direct Sanger Sequencing

For *swrB* point mutations generated through PCR mutagenesis, a PCR product containing the *swrB* open reading frame was amplified from *B. subtilis* chromosomal DNA (either from strain 3610 or the appropriate mutant strain) using the primer set 2429/2430. The *swrB* PCR product was then sequenced using primer 2429 and 2430 individually.

### SPP1 phage transduction

To 0.2 ml of dense culture grown in TY broth (LB broth supplemented after autoclaving with 10 mM MgSO_4_ and 100 µM MnSO_4_), serial dilutions of SPP1 phage stock were added and statically incubated for 15 minutes at 37°C. To each mixture, 3 ml TYSA (molten TY supplemented with 0.5% agar) was added, poured atop fresh TY plates, and incubated at 30°C overnight. Top agar from the plate containing near confluent plaques was harvested by scraping into a 15 ml conical tube, vortexed, and centrifuged at 5,000 x g for 5 minutes. The supernatant was treated with 25 µg/ml DNase final concentration before being passed through a 0.45 µm syringe filter and stored at 4°C.

Recipient cells were grown to stationary phase in 3 ml TY broth at 37°C. 1 ml cells were mixed with 25 µl of SPP1 donor phage stock. 9 ml of TY broth was added to the mixture and allowed to stand at 37°C for 30 minutes. The transduction mixture was then centrifuged at 5,000 x g for 5 minutes, the supernatant was discarded and the pellet was resuspended in the remaining volume. 100 µl of cell suspension was then plated on LB fortified with 1.5% agar, the appropriate antibiotic, and 10 mM sodium citrate.

### Swarm expansion assay

Cells were grown to mid-log phase at 37°C in LB broth and resuspended to 10 OD_600_ in pH 8.0 PBS buffer (137 mM NaCl, 2.7 mM KCl, 10 mM Na_2_HPO_4_, and 2 mM KH_2_PO_4_) containing 0.5% India ink (Higgins). Freshly prepared LB containing 0.7% Bacto agar (25 ml/plate) was dried for 10 minutes in a laminar flow hood, centrally inoculated with 10 µl of the cell suspension, dried for another 10 minutes, and incubated at 37°C. The India ink demarks the origin of the colony and the swarm radius was measured relative to the origin. For consistency, an axis was drawn on the back of the plate and swarm radii measurements were taken along this transect. For experiments including IPTG, cells were propagated in broth in the presence of IPTG, and IPTG was included in the swarm agar plates.

### Plate-based PhoA and LacZ assays

Stocks 5-bromo-4-chloro-3-indolyl-β-D-galactopyranoside (X-Gal, Sigma-Aldritch) and 5-bromo-4-chloro-3-indolyl-phosphate disodium salt (X-P, Sigma-Aldritch) were made immediately before use by dissolving 20 mg of either compound in 1ml dimethylformamide (DMF) or 1 ml deionized water, respectively. The X-P solution was filter sterilized and both solutions were stored in opaque Eppendorf tubes. 100 ul of each solution was spread atop separate LB plates fortified with 1.5% agar containing 1 mM IPTG and the solutions were allowed to absorb into the agar for 1 hr at 37°C in the dark. Cells were then struck on the plates and incubated at 30°C overnight. The back slides of the Petri plates were photographed using a Kodak Pixpro FZ53 digital camera.

### Western blotting

*B. subtilis* strains were grown in LB broth to OD_600_ ~0.5, 1 ml was harvested by centrifugation, and resuspended to 10 OD_600_ in Lysis buffer (20 mM Tris pH 7.0, 10 mM EDTA, 1 mg/ml lysozyme, 10 μg/ml DNAse I, 100 μg/ml RNAse I, 1 mM PMSF) and incubated 30 minutes at 37°C. Each lysate was then mixed with the appropriate amount of 6x SDS loading dye to dilute the loading dye to 1x concentration. Samples were separated by 12% Sodium dodecyl sulfate-polyacrylamide gel electrophoresis (SDS-PAGE). The proteins were electroblotted onto nitrocellulose and developed with a 1:10,000 dilution of (anti-SwrB), 1:10,000 dilution of (anti-GFP), or 1:80,000 dilution of (anti-SigA) of primary antibody and a 1:10,000 dilution secondary antibody (horseradish peroxidase-conjugated goat anti-rabbit immunoglobulin G). Immunoblot was developed using the Immun-Star HRP developer kit (Bio-Rad).

### Microscopy

Fluorescence microscopy was performed with a Nikon 80i microscope along with a phase contrast objective Nikon Plan Apo 100X and an Excite 120 metal halide lamp. Alexa Fluor 594 C_5_ maleimide fluorescent signals were visualized with a C-FL HYQ Texas Red Filter Cube (excitation filter 532-587 nm, barrier filter >590 nm). GFP was visualized using a C-FL HYQ FITC Filter Cube (FITC, excitation filter 460-500 nm, barrier filter 515-550 nm). YFP was visualized using a C_FL HYQ YFP Filter Cube (excitation filter 490-510 nm, barrier filter 515-550 nm). TMA-DPH fluorescent signal was visualized using a UB-2E/C DAPI Filter Cube (excitation filter 340-380 nm, barrier filter 435-485 nm). Images were captured with a Photometrics Coolsnap HQ^2^ camera in black and white, false colored and superimposed using Metamorph image software.

For *P*_*flache*_*-swrB-YFP* microscopy, cells were grown at 37°C in LB broth to OD_600_ 0.6-1.0, resuspended in 30 μl PBS buffer containing 0.1mM TMA-DPH and incubated for 5 min at room temperature. The cells were pelleted, resuspended in 30μl PBS buffer, and were observed by spotting 4 μl of suspension on a cleaned microscope slide and immobilized with a poly-L-lysine-treated glass coverslip.

For fluorescent microscopy of *swrBΩGFP* and flagellar hooks, 1.0 ml of broth culture was harvested at 0.6-1.0 OD_600_, resuspended in 50 μl of PBS buffer containing 5μg/ml Alexa Fluor 594 C_5_ maleimide (Molecular Probes), incubated for 3 min at room temperature, and washed once in 1.0 ml of PBS buffer. The suspension was pelleted, resuspended in 30 μl of PBS buffer containing 0.1mM TMA-DPH, and incubated for 5 min at room temperature. The cells were pelleted, resuspended in 30 μl PBS buffer, and were observed by spotting 4 μl of suspension on a cleaned microscope slide and immobilized with a poly-L-lysine-treated glass coverslip.

### β-Galactosidase assays

Cells were harvested from cultures growing at 37°C in LB broth. Cells were collected in 1.0 ml aliquots and suspended in an equal volume of Z buffer (40 mM NaH_2_PO_4_, 60 mM NaHPO_4_, 1.0 mM MgSO_4_, 10 mM KCl, and 38 mM 2-mercaptoethanol). Lysozyme was added to each sample to a final concentration of 0.2 mg/ml and incubated at 37°C for 30 min. Each sample was diluted in Z buffer to a final volume of 500 μl, and the reaction was started with 100 μl of 4 mg/ml 2-nitrophenyl β-galactopyranoside (ONPG, Sigma-Aldritch) in Z buffer and stopped with 250 μl of 1M Na_2_CO_3_. The OD_420_ of the reaction mixture was measured, and the β-galactosidase-specific activity was calculated according the equation [OD_420_/(time X OD_600_)] X dilution factor X 1000.

## ACKNOWLEDGEMENTS

We are grateful to Cristina Landeta for guidance in the development of the alkaline phosphatase reporter system. Special thanks go to Kearns lab members for their intellectual support; Loralyn Cozy, Joyce Patrick, Rebecca Calvo, Melissa Konkol, Eric Vanderpool, Anna Bree, Sampriti Mukherjee, and others. This work was funded by USDA grant WIS02030 to BMB and National Institutes of Health R35 grant GM131783 to DBK.

